# Global connectivity fingerprints predict the domain generality of multiple-demand regions

**DOI:** 10.1101/2021.01.25.428141

**Authors:** Douglas H. Schultz, Takuya Ito, Michael W. Cole

## Abstract

A set of distributed cognitive control networks are known to contribute to diverse cognitive demands, yet it is unclear how these networks gain this domain-general capacity. We hypothesized that this capacity is largely due to the particular organization of the human brain’s intrinsic network architecture. Specifically, we tested the possibility that each brain region’s domain generality is reflected in its level of global (hub-like) intrinsic connectivity, as well as its particular global connectivity pattern (*connectivity fingerprint*). Consistent with prior work, we found that cognitive control networks exhibited domain generality, as they represented diverse task context information covering sensory, motor response, and logic rule domains. Supporting our hypothesis, we found that the level of global intrinsic connectivity (estimated with resting-state fMRI) was correlated with domain generality during tasks. Further, using a novel information fingerprint mapping approach, we found that each cognitive control region’s unique rule response profile (*information fingerprint*) could be predicted based on its unique intrinsic connectivity fingerprint and the information content in non-cognitive control regions. Together these results suggest that the human brain’s intrinsic network architecture supports its ability to represent diverse cognitive task information, largely via the location of multiple-demand regions within the brain’s global network organization.

## Introduction

Cognitive control processes allow individuals to intentionally select thoughts and behaviors according to current goals. These processes are particularly important in situations when previously learned rules need to be applied in novel situations. This ability to transfer rules from one context and apply them in a new situation greatly increases cognitive flexibility. Rapid instructed task learning (RITL) is a cognitive ability that many, including ourselves, have used to study cognitive control that requires especially flexible cognitive processes (Cole, Laurent, et al., 2013; Liefooghe et al., 2013; Pereg & Meiran, 2019; Ruge et al., 2019). These cognitive control processes are supported by specialized large-scale brain networks (Cole & Schneider, 2007; Niendam et al., 2012), which are thought to contribute to the ability to direct cognition in a goal-directed manner by flexibly reconfiguring domain-specific (e.g., sensory and motor) neural systems (Cole et al., 2017; Cole, Reynolds, et al., 2013; Desimone & Duncan, 1995; Miller & Cohen, 2001; Schneider, 2003).

Historically, there has been a debate between theories emphasizing the localization of brain function and those that emphasize a highly distributed organization in the brain. At the spatial scale of fMRI, evidence for cognitive control networks suggests many cognitive processes are neither fully distributed nor fully localized to a single brain region, since it appears cognitive control processes are distributed across regions within these brain networks. The proposal for a multiple demand network followed the observation of common patterns of activation during a variety of cognitive tasks including: auditory discrimination, visual divided attention, self-paced response production, task switching, spatial problem solving, and semantic processing of words (Duncan, 2010; Duncan & Owen, 2000). These findings suggest that regions belonging to the multiple demand network are recruited under a large variety of cognitive processes, regardless of the modality of the stimuli or specific cognitive demand. Despite the common patterns of co-activation observed in the multiple demand network, subsequent research has suggested that this network is not homogenous (Camilleri et al., 2018). The multiple demand network can be split into several smaller cognitive control networks characterized by different cognitive specializations (Assem et al., 2020; Cole & Schneider, 2007; Yeo et al., 2015). This suggests that task-relevant representations show different patterns of localization, and that they aren’t homogeneously distributed across regions composing the multiple demand network.

Going beyond task-evoked activity amplitudes, analyses of multivariate task activation patterns has provided important insights into the information content contained within cognitive control networks. For instance, a number of studies have found that rule identity can be decoded based on multivariate activation patterns, and that the most accurate decoding of these patterns tends to occur in cognitive control network regions (Cole et al., 2011; Pischedda et al., 2017; Reverberi, Görgen, et al., 2012; Reverberi, Gorgen, et al., 2012; Woolgar et al., 2011; Yeo et al., 2015). Electrophysiological studies in monkeys have also identified neurons that code for specific task rules, largely in prefrontal cortical regions (Asaad et al., 2000; Brincat et al., 2018; Wallis & Miller, 2003; White & Wise, 1999). These context-dependent neural representations within cognitive control networks are likely crucial for increasing cognitive flexibility during goal-directed cognitive tasks.

We previously found that multivariate patterns of activation within cognitive control networks contain information across three distinct cognitive domains (logic, sensory, and motor rules) in the concrete permuted rule operations (C-PRO) paradigm. The C-PRO paradigm consists of a large set of systematically related flexible control tasks (Cocuzza et al., 2020; Ito et al., 2017). We sought to replicate these results and expand on them by testing hypotheses regarding how the brain’s intrinsic network architecture contributes to diverse task rule representations. Previous studies have found support for both distributed and localized task rule representations in the human brain. This may reflect the possibility that task rule representation occurs on a continuum, from highly distributed to largely localized. We predicted that cognitive control brain regions would contain task rule representations, or task information, across a wide variety of rule domains, consistent with previous research supporting distributed rule representations across the multiple demand network. We also expected to observe a degree of localization in task rule representations. For example, we predicted task rule information specific to the motor rules to be largely localized to the motor cortex. While there is evidence for both localized and distributed rule representations in the brain, this study was designed to evaluate the degree of specialization of task rule representations.

While there is a great deal of evidence supporting the existence of a multiple-demand network, it is unclear how this domain generality emerges (Assem et al., 2020). Building on our prior work demonstrating that regions within this network are hubs with especially widespread intrinsic connectivity (Cocuzza et al., 2020; Cole, Bagic, et al., 2010; Cole et al., 2012; Schultz et al., 2018), we hypothesized that domain generality may emerge from the widespread hub connectivity of these brain regions. However, not all cognitive control regions are equally domain general (Assem et al., 2020), suggesting a role for the particular intrinsic connectivity patterns (the *connectivity fingerprints*) of these cognitive control regions in shaping their specific level of domain generality. We recently found that localized task-evoked responses can be accurately predicted by distributed connectivity-based activity flow simulations (which model the movement of task-evoked brain activity between brain regions), suggesting prominent roles for distributed network-level processes in determining many localized cognitive functions (Cole et al., 2016; Hearne et al., 2021). Based on these and related results, we developed the more specific hypothesis that patterns of information content (*information fingerprints*) in cognitive control regions are determined by (and can therefore be predicted by) information content in other brain regions weighted by their connectivity to cognitive control regions. If task rule representations in cognitive control networks can be predicted based on global intrinsic functional connectivity patterns, it would further support the hypothesis that global connectivity plays an important role in task rule representation. Alternatively, the intrinsic functional connectivity patterns may not be important to task rule representation. For example, the internal processing of information within a functional node may be the critical feature determining task rule representation with connectivity patterns being less important. However, if the intrinsic network architecture is important, we would expect domain generality in cognitive control networks to be predictable based on the specific global connectivity profiles of regions within these networks. Determining how these domain-general representations are generated will be important, given that they may serve to integrate domain-specific representations so they can be coordinated for task performance when different combinations of rules are encountered, likely increasing the cognitive flexibility of the human brain.

## Methods

### Participants

Data were collected at the Rutgers University Brain Imaging Center (RUBIC). The participants were recruited from the Rutgers University-Newark campus and surrounding community. All procedures were approved by the Rutgers University-Newark Institutional Review Board and all participants provided informed consent. The sample consisted of data from 106 participants. Technical error or equipment malfunction during the scanning session resulted in removing six participants from the study. Out of the remaining 100 participants, 56 were female and 44 were male. Participants were between the ages of 18 and 38 (*M* = 22.24, *SD* = 4.07). Participants were all right-handed, had not been diagnosed with any psychiatric disorders, and met standard MRI safety criteria.

### Behavioral Task

We used the C-PRO paradigm (Ito et al., 2017), which is a variant of the original permuted rule operations task (Cole, Bagic, et al., 2010). The C-PRO task combines specific task rules from three different rule domains (logical, sensory, and motor) to create sixty-four unique task sets (Figure 1). Each of the three different rule domains includes four specific rules (Logic: Both, Not Both, Either, Neither; Sensory: Red, Vertical, Constant, High Pitch; Motor: Left Index, Left Middle, Right Index, Right Middle). This task design allowed us to compare task rule information content across three different modalities while controlling for attention, arousal, sensory input, and motor output across tasks. Visual and auditory stimuli were presented simultaneously on each trial with the visual stimuli consisting of red or blue bars that were oriented either horizontal or vertical. The auditory stimuli were either high (3000 Hz) or low (300 Hz) tones that were presented continuously or non-continuously (beeping). Trials were structured as miniblocks. Each miniblock started by presenting instructions for 3925 ms. After the instructions there was a jittered delay that lasted between 1570 and 6280 ms. Three trials were presented. The duration of each trial was 2355 ms. The intertrial interval was 1570 ms. Following the last trial there was another jittered delay between 7850 and 12560 ms. The mean miniblock duration was 28,260 ms. Each of the sixty-four task set miniblocks were presented twice during the experiment, and the same task set was never presented consecutively.

**Figure 1.**
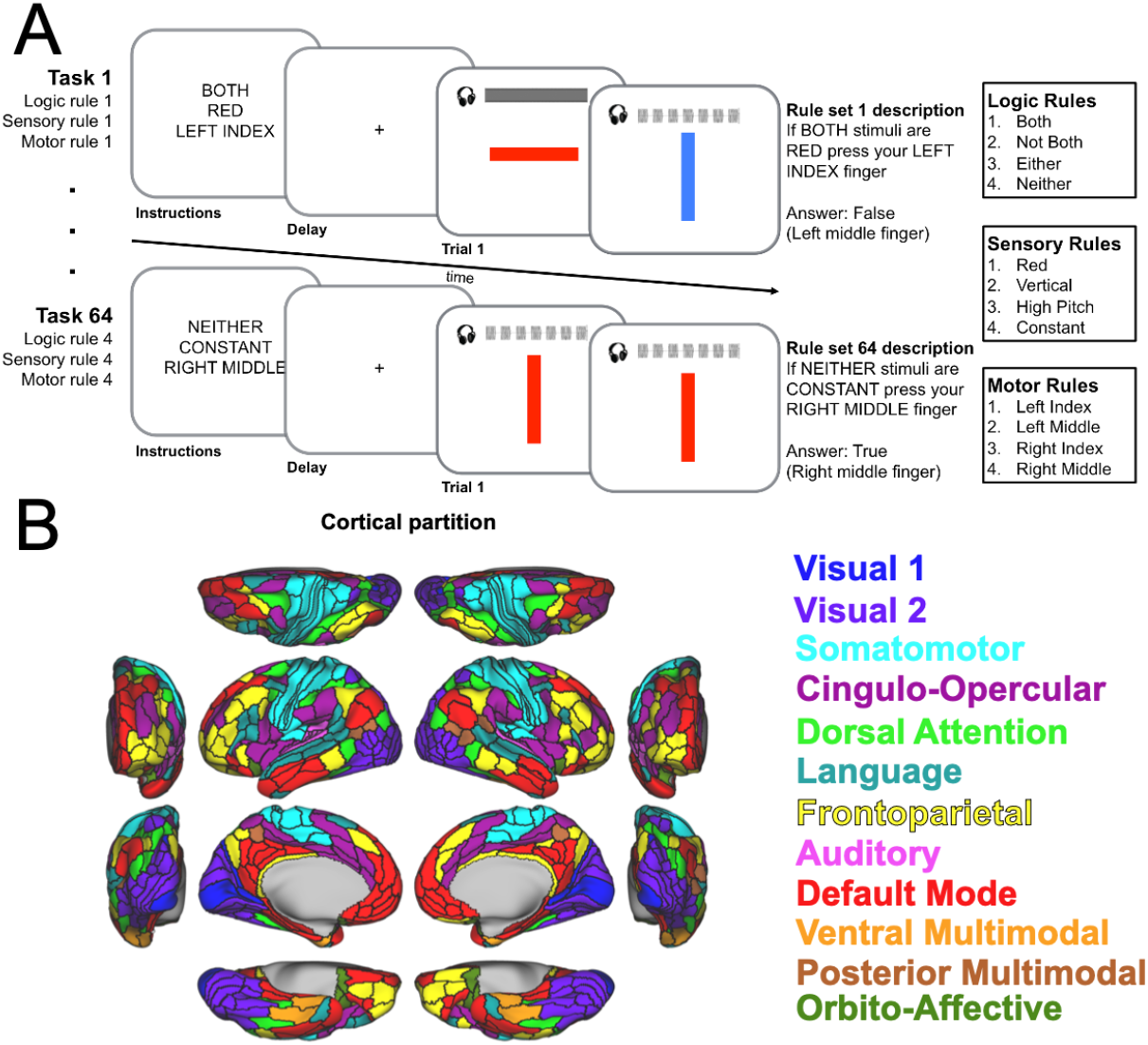
The Concrete Permuted Rule Operations cognitive paradigm and functional network assignments. (A) The cognitive paradigm combines specific task rules from three different rule domains (logical, sensory, and motor) to create sixty-four unique task sets. (B) Functional network assignment for brain regions based on (Ji et al., 2019).

### MRI Parameters

Multiband whole-brain echo-planar imaging (EPI) data was collected using a 32-channel head coil on a 3T Siemens Trio MRI scanner with the following parameters: TR = 785 ms, TE = 34.8 ms, flip angle = 55°, Bandwidth 1924/Hz/Px, in-plane FoV read = 208 mm, 72 slices, 2.0 mm isotropic voxels, with a multiband acceleration factor of 8. Whole-brain high-resolution T1-weighted and T2-weighted anatomical scans with 0.8 mm isotropic voxels were also collected. Spin echo field maps were collected in both the anterior to posterior direction and the posterior to anterior direction consistent with the Human Connectome Project preprocessing pipelines (Glasser et al., 2013). Resting state fMRI was collected prior to the task fMRI scans as described previously (Schultz et al., 2018). The resting state scan was 14 minutes in duration (1070 TRs). Eight runs of task fMRI data were collected while participants performed the C-PRO task. Each task run had a duration of 7 minutes and 36 seconds (581 TRs). Task runs were run sequentially with a short break between each run.

### fMRI Preprocessing

We minimally preprocessed the fMRI data using the publicly available Human Connectome Project minimal preprocessing pipeline (version 3.5.0). This included anatomical reconstruction and segmentation, EPI reconstruction, segmentation, normalization to a standard template, intensity normalization, and motion correction (Glasser et al., 2013). The transformation of volume data to the cortical surface is part of this pipeline. Subsequent processing was conducted on CIFTI 64k gray ordinate space. A standard general linear model (GLM) was fit to the vertex-level task time series with a convolved canonical hemodynamic response function from SPM using the sixty-four task sets as regressors. Task regressors represented the entire duration of each miniblock. We also included 12 motion parameters (6 motion estimates and their derivatives), mean signal from the ventricles and white matter (and their derivatives) which were defined anatomically via Freesurfer (Fischl et al., 2002) as nuisance regressors. Motion scrubbing as described by Power and colleagues (2012) was implemented with a frame-wise displacement threshold of 0.3 mm. Frame-wise displacement estimates were temporally filtered prior to thresholding to reduce the influence of respiration on FD estimates (Siegel et al., 2016). Resting state data was processed in the same way as task data with the exception of excluding the task based regressors. Functional connectivity was estimated by calculating Pearson’s correlation of the mean timeseries for each pair of regions in the brain. Brain regions were defined by the Glasser et al. (2016) parcellation which consists of 360 cortical regions. Brain regions were assigned to functional networks as in Ji and colleagues, (2019).

### fMRI Data Analysis

#### Information estimates

Information estimates were computed for each brain region (see Ito et al., 2017). The calculation of information estimates was conducted within each participant using a cross-validated multivariate pattern analysis.

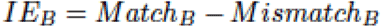

Task beta estimates were created for each vertex and for each of the sixty-four unique task sets. A representational analysis using a minimum-distance metric (using Spearman correlation as the distance measure) was then calculated for each of the 360 cortical regions. Information estimates were calculated on multiple levels (Mur et al., 2009). First, we calculated information estimates for each domain (Logic, Sensory, Motor). For each of the sixty-four task sets we calculated the similarity between the vertex-level pattern of activation (Bk) with the vertex-level pattern of activation in the remaining sixty-three task sets that shared the rule of the domain being examined (Bmatch) with a Spearman correlation. There were fifteen other task sets that shared the rule of the domain being examined (K).

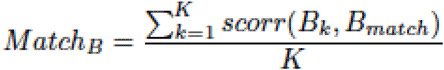

Next we calculated the similarity between the vertex-level pattern of activation (Bk) with the vertex-level pattern of activation for each of the sixty-three remaining tasks which did not share the rule of the domain being examined (we excluded the task sets that shared off-domain rules with the held out task set, 45 task sets (N)).

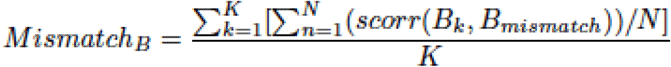

We calculated the mean similarity for task sets with shared rules and the mean similarity for task sets without a shared rule in the same domain. The mean of these values across the sixty-four task sets was calculated. Then we subtracted the non-shared rule similarity from the shared rule similarity (Figure 2). Thus, positive information estimates indicate that the vertex-wise pattern of activation is more similar to the vertex-wise pattern of activation for trials in which shared rules for each domain were present relative to the pattern of activation for trials in which non-shared rules for each domain were present. This results in an information estimate for each domain (Logic, Sensory, Motor) for each brain region. We used a similar procedure to calculate information estimates for each individual task rule. Rather than comparing all task sets that shared any rule in a particular domain, we calculated the information estimates for only task sets that shared a specific rule in each domain. This process results in an information estimate for each individual task rule for each brain region (Logic: Both, Not Both, Either, Neither; Sensory: Red, Vertical, Constant, High Pitch; Motor: Left Index, Left Middle, Right Index, Right Middle). Statistical comparisons were made for each region by using a *t*-test against zero. The p-values from these tests were corrected for multiple comparisons using permutation tests where condition labels were shuffled (Nichols & Holmes, 2002) with 1000 permutations unless otherwise specified, and a FWE corrected threshold of p < 0.05 was used.

**Figure 2.**
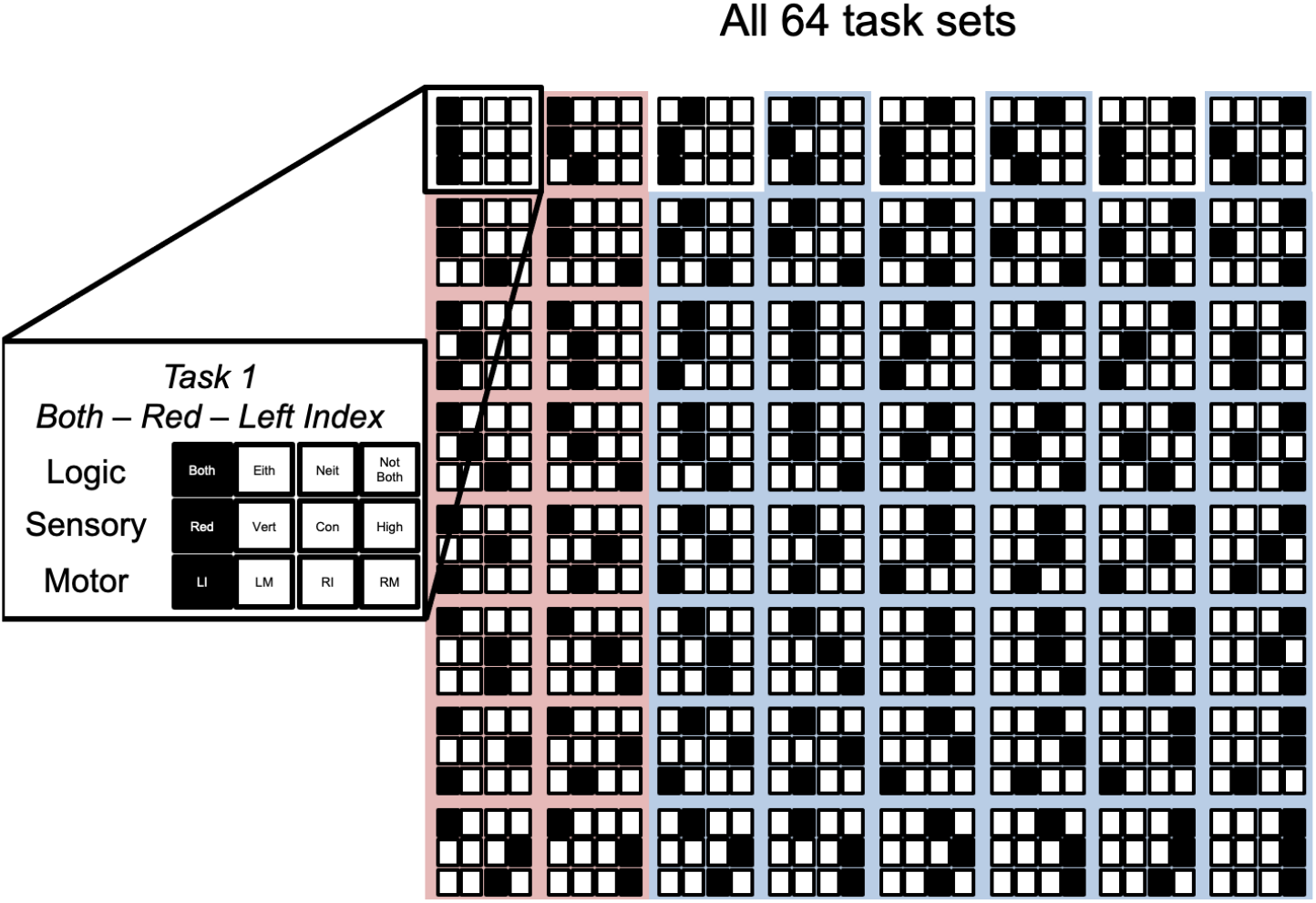
Calculation of information estimates. For each task set (Task 1 in this example), we calculated the similarity of the vertex-wise activation pattern for this task set with other task sets that shared a rule (those in the red area for the Both rule in this example) and other task sets that did not share that rule (those in the blue area for the Both rule in this example).

#### Information fingerprint mapping

Information fingerprints were defined for each brain region as the pattern of the information estimates for each of the 12 task rules (Logic: Both, Not Both, Either, Neither; Sensory: Red, Vertical, Constant, High Pitch; Motor: Left Index, Left Middle, Right Index, Right Middle). We used a modified version of activity flow mapping (Cole et al., 2016) to predict information fingerprints. For each participant we predicted the information estimate for each task rule by weighting the information estimate for every other region in the brain by the strength of the functional connectivity between the two regions. Importantly, the functional connectivity estimates were based on an independent resting state scan. After calculating the predicted information estimates, we combined the 12 individual task rule information estimates into information fingerprints and correlated the predicted information fingerprints with the actual information fingerprints. The resulting Pearson’s r values were Fisher’s Z transformed, and group statistical comparisons were made using a *t*-test against zero. The p-values from these tests were corrected for multiple comparisons using permutation tests where condition labels were shuffled 1000 times unless specified otherwise, and a FWE corrected threshold of p < 0.05 was used.

## Results

### Participants learned the C-PRO paradigm

Cross-subject average accuracy on the C-PRO task was 84.86%. Participants performed well above chance level performance (25%) on all 12 rules: Both (*M(SD)*, 91.2%(0.07%)); Either (89.1(0.09)); Neither (81.4(0.11)); Not Both (72.6(0.14)); Constant (78.2(0.15)); High Pitch (83.8(0.10)); Red (86.7(0.08)); Vertical (85.7(0.09)); Left Index (83.2(0.10)); Left Middle (84.0(0.10)); Right Index (83.5(0.10)); Right Middle (83.6(0.10)). No participants performed at less than chance level on any of the rules.

### Task domain information is present in numerous brain regions

We tested whether the activation pattern within any brain region contained information about each of the three task domains (Logic, Sensory, Motor). This allowed us to map task rule information and determine the degree of distribution for each rule. We calculated the information estimate for each region on each of the task domains. It is also important to note that the task was counterbalanced in order to ensure that there was no systematic difference in what stimuli were presented to participants across rules so that task activation related to the auditory and visual stimuli presented did not bias information estimates.

The information estimate measure has previously been validated using a computational model and a portion of the same data analyzed here (Ito et al., 2017). Briefly, the information estimate compares the similarity in activation pattern (Spearman correlation) for a given task (e.g., Both, Red, Left Index) with the activation pattern from other tasks with a matching rule. That value is compared to the similarity in activation pattern between the same starting task (Both, Red, Left Index) and the activation pattern from the other tasks that do not share the same rule for each participant. An information estimate value greater than zero suggests that the pattern of activity for a particular rule is more similar to tasks that share a rule than it is for tasks that do not share a rule. If the information estimate value is significantly greater than zero we conclude that the activation pattern in that particular brain region contains information related to the rule.

First we tested whether brain regions contained general information about the three different modalities (Logic, Sensory, Motor). For example, a region contained information regarding the Logic rule domain if the pattern of activity was more similar on tasks that shared any Logic rule. We found that widespread regions of the brain contained information regarding the Logic and Sensory rules. We found that the brain regions containing information about the Motor rules were largely located in the somatomotor network, with several adjacent regions from other networks containing information (Figure 3).

**Figure 3.**
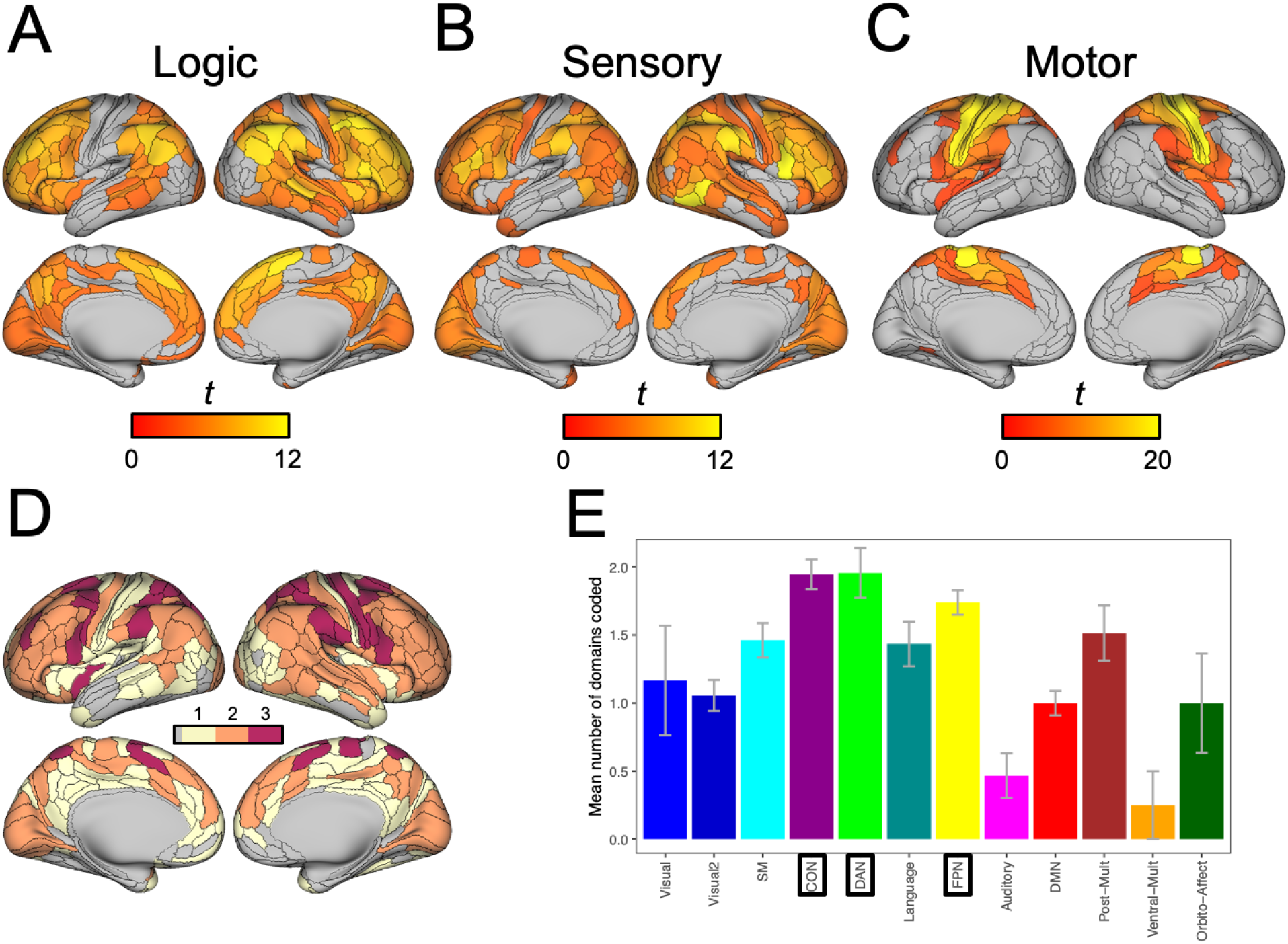
Brain regions contain information across content domains. Vertex activity within each brain region can be used to decode rules. (A) Widespread cortical regions from many networks contain information for the logic rules. Only significant results are shown. Non-significant results appear as gray for panels A, B, and C. (B) Significant sensory rule information is also evident in a large number of regions and many networks. (C) Sensorimotor network brain regions contain the most information for the motor rules. The degree of information content in the sensorimotor network is also greater than that observed in either the logic or sensory rules. (D) Conjunction between panels A, B, and C showing how many domains each brain region represents. (E) Summarizing the mean number of domains that are represented in each functional network. Cognitive control networks (marked by black boxes) contain information about the highest number of domains. Error bars indicate standard error.

One aspect of the task that could not be addressed with counterbalancing was the possible influence of activity related to motor responses on information estimates. For each motor rule (Left Index, Left Middle, Right Index, Right Middle), participants were much more likely to make a motor response with the hand specified in the rule rather than responses on the other hand. To address this issue, we ran another version of the GLM that included four regressors – one for each type of motor response participants made. Given that these motor response regressors differed from the motor rule regressors (which were of interest), the motor response regressors were treated as nuisance regressors and we repeated the information estimate calculations. We found that the magnitude of the results for the Motor rule domain decreased (largest *t* in the original analysis = 20.44, largest *t* after including the motor responses in the GLM = 18.07). The magnitude of the effect in the Logic and Sensory domains were unchanged (Logic: largest *t* in the original analysis = 11.59, largest *t* after including the motor responses in the GLM = 11.57; Sensory: largest *t* in the original analysis = 13.28, largest *t* after including the motor responses in the GLM = 13.32). Additionally, we examined the pattern of information estimates across the brain for each domain. The pattern of results in the Motor rule domain was very similar in the original analysis and in the analysis including the motor response regressors (Spearman’s rho = 0.707, p < 0.0001). The similarity in the results for the Logic and Sensory domain was even higher (Logic: rho = 0.977, Sensory: rho = 0.986). Together, these results suggest that activity related to motor responses is not systematically biasing information estimates.

Next, we used the same process to evaluate whether any brain regions contained information for each of the twelve individual rules (Both, Not Both, Either, Neither, High, Constant, Red, Vertical, Left Index, Left Middle, Right Index, Right Middle). For example, the similarity in the pattern of activation between a particular task (Both, Red, Left Index) and other tasks containing the Both rule relative to tasks containing other Logic rules. This resulted in twelve FWE-corrected maps, one for each individual rule (Figure 4).

**Figure 4.**
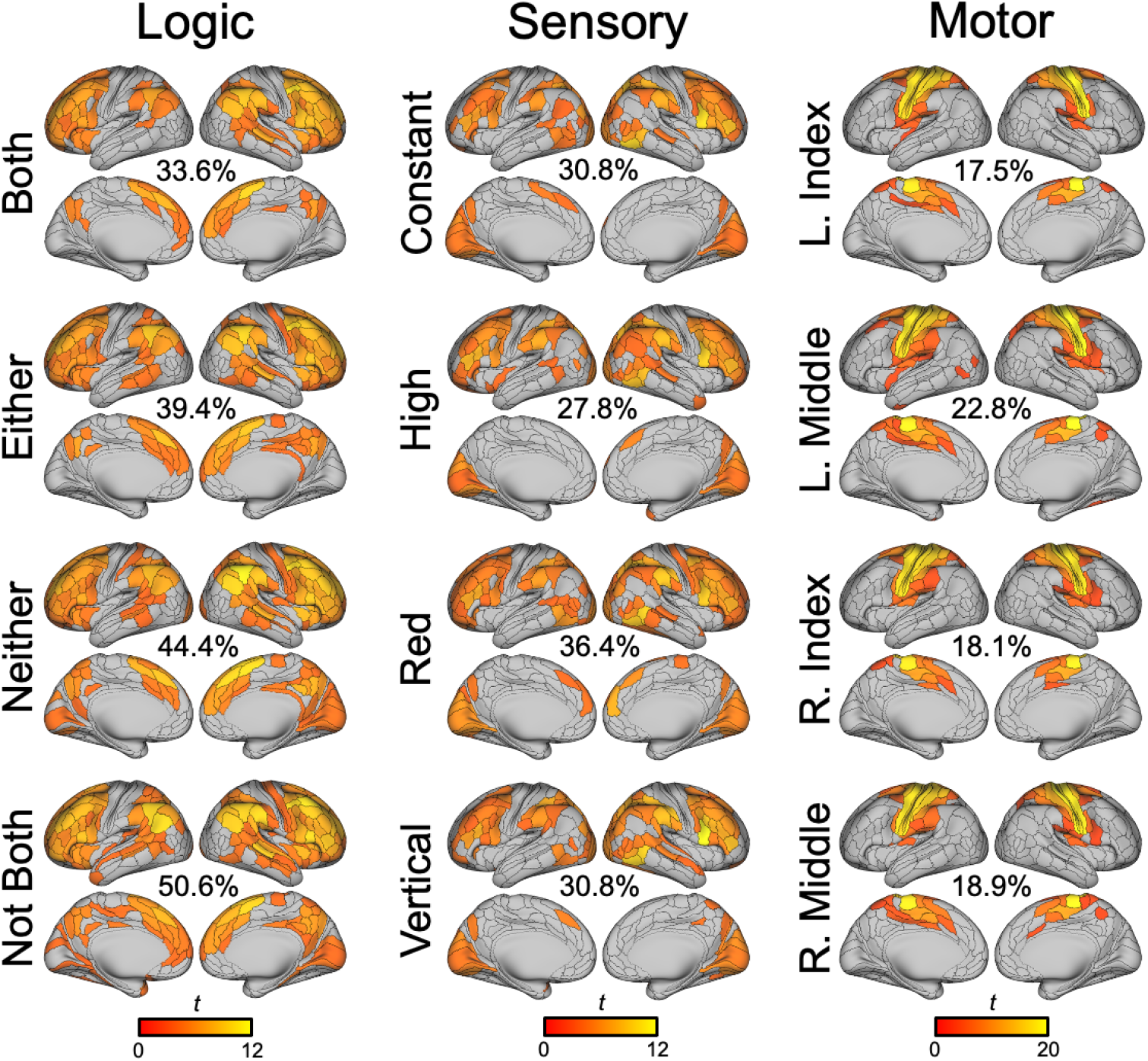
Information estimates for each of the individual rules are largely consistent within each modality. Significant information estimates (FWE corrected p < 0.05) are shown for each of the twelve rules. Each column represents a rule domain and rows are the four rule variants within each domain. The information estimates for each individual rule are also consistent with the more general information estimates calculated at the domain level. The percentage of cortical regions (out of 360) showing significant information estimates is provided for each rule.

### Cognitive control networks contain domain general information

Next, we assessed whether any brain regions contained domain general information. The brain maps for each individual rule (Figure 4) were binarized, and if a brain region contained a significant degree of information (FWE-corrected) it was coded as a 1. If a brain region did not contain a significant degree of information it was coded as a 0. Then we determined the mean number of rules that were coded for each domain in each brain region. The mean across the three domains was calculated for each region providing an estimate of the degree of domain generality. A truly domain general brain region would contain information for all rules across all domains. Based on our metric this brain region would get a score of 4. A brain region that contained information about all four rules in one domain, but no rules in the other two domains would get a score of 1.33. We found that cognitive control networks (DAN, FPN, CON) had the largest values suggesting that they contained information from a wide variety of rules and domains (Figure 5). The cognitive control networks coded for a larger number of rules per domain than other brain regions, *t*(358) = 3.61, p = 0.00035.

**Figure 5.**
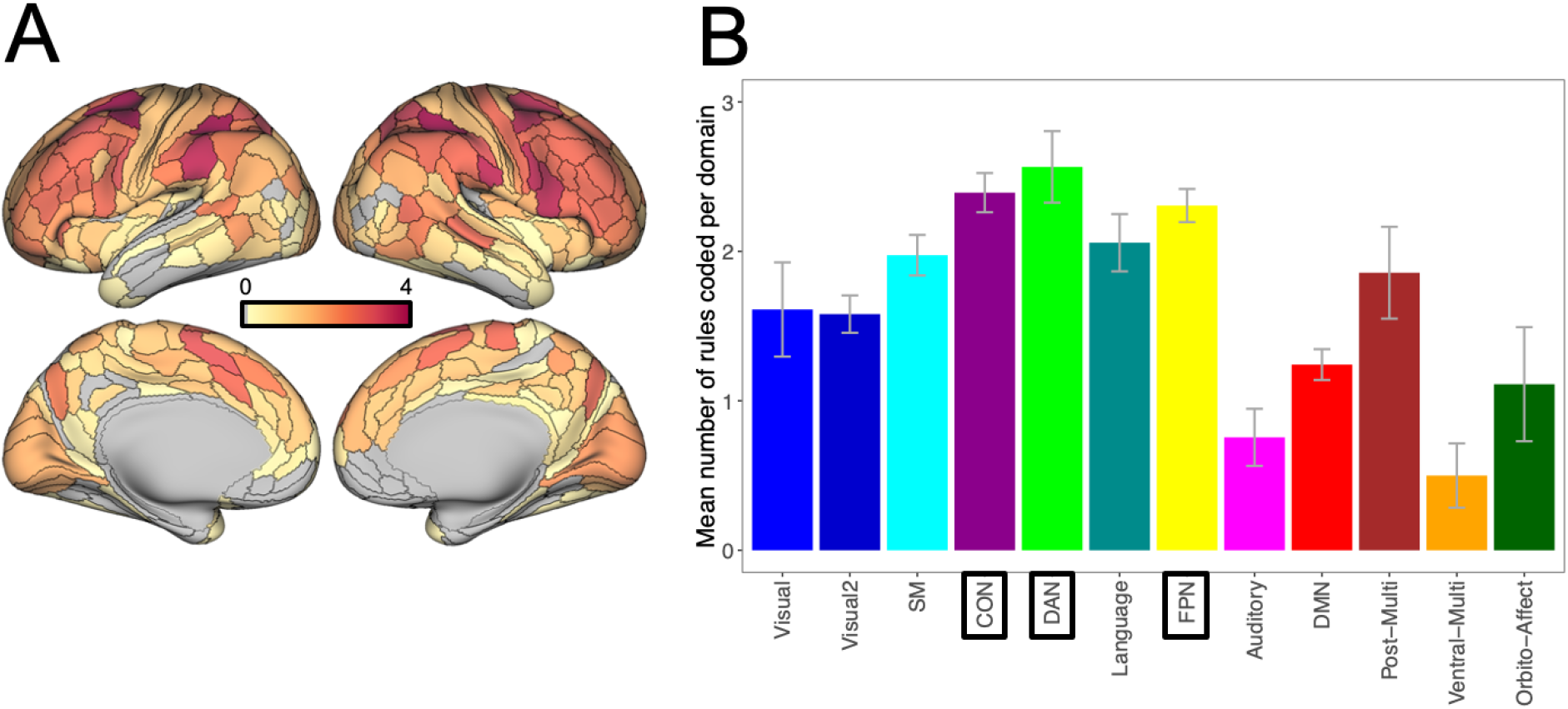
Cognitive control networks show domain general properties. (A) The mean number of rules coded across domains for each brain region. Brain regions colored in red contain information for all four rules in each domain. Brain regions depicted in white may only code for one specific rule in one particular domain. (B) Summarizing the mean number of rules coded per domain across the functional networks reveals that cognitive control networks (marked by black boxes) are domain general meaning they contain information for a larger number of rules than other brain networks. Error bars indicate standard error.

### Mapping domain specific information

We next sought to contrast with domain generality, identifying brain regions that preferentially contained information for a specific rule modality. For each rule modality we identified brain regions that contained information for a larger number of rules than the remaining modalities. Within these regions we calculated the difference between the number of rules coded for the modality of interest and subtracted the mean number of rules coded of the remaining modalities. A larger number would indicate a greater imbalance in task rule information suggesting that the region preferentially contained task rule information pertaining to the modality of interest (Figure 6). Qualitatively, we found that the DMN contained domain specific information related to the Logic rules. We also observed that the visual and auditory networks contained domain specific information related to the Sensory rules. Additionally, the somatomotor network appeared to contain domain specific information related to the Motor rules.

**Figure 6.**
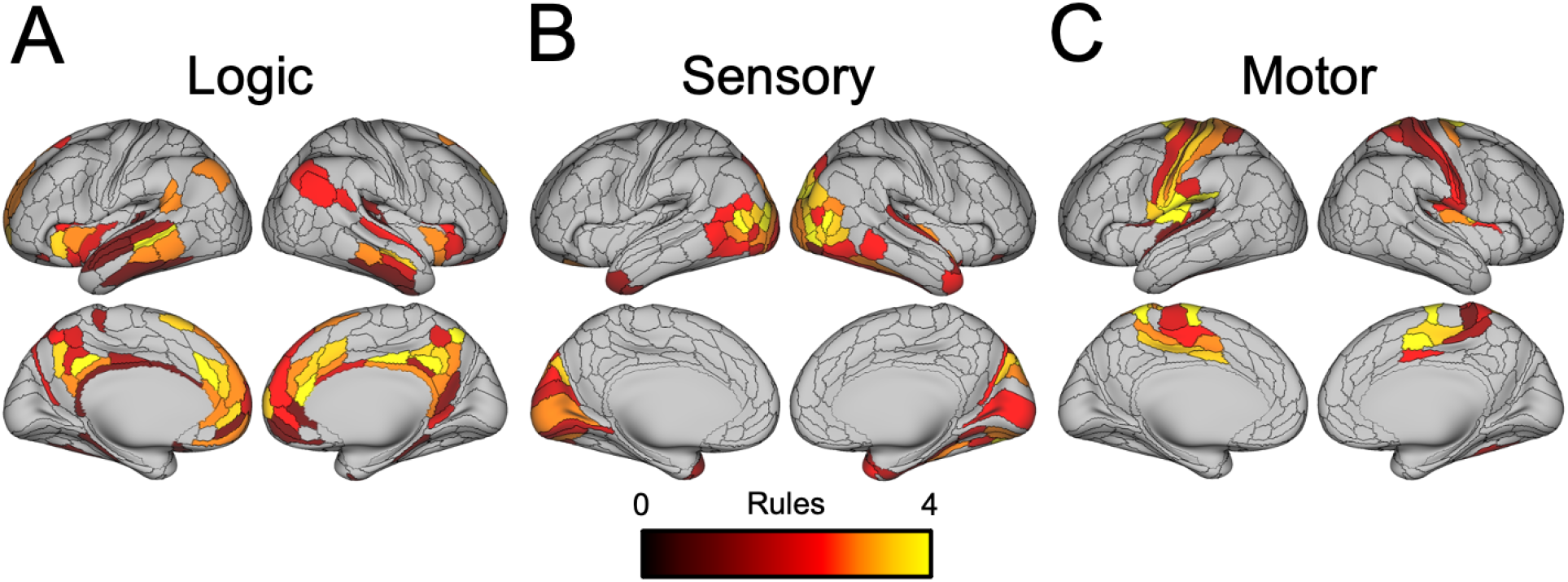
Some regions contain domain specific information. Panels reflect the difference of rules coded for each domain from the mean number of rules coded for the other two domains. (A) Some regions contain more information for the logic domain than the sensory and motor domains. Many of the regions containing logic domain specific information are in the DMN. (B) Some regions contain more information for the sensory rules than the logic and motor rules. The stimuli presented were visual and auditory. Domain specific information for the sensory rules is primarily located in the visual and auditory networks. (C) Some regions contain more information for the motor rules than the logic and sensory rules. Domain specificity for the motor rules is primarily observed in the somatomotor network.

### Domain general regions are active during many task contexts and are more hub-like

We ran a standard GLM using 64 task regressors (one for each possible rule combination) and tested which brain regions showed a significant (FWE-corrected) level of activation for each of the 64 unique rule combinations. Then we calculated a summary statistic for each region reflecting what percentage of the 64 tasks showed significant positive activation (Figure 7A). It is important to note that the results of this task activation analysis are not only sensitive to activity related to each of the unique rule combinations, but also to shared cognitive processes that are common to the general structure of the task such as reconfiguration, task switching, and working memory. Finally, we compared the percentage of activation for the unique rule combinations in domain general brain regions (mean rules coded per domain > 3) and the rest of the brain. The probability of a brain area showing significant activity during the task was not normally distributed so we used the Wilcoxon signed rank test to evaluate the difference in the probability of activation during the task for domain general regions and the rest of the brain. Domain general regions are more likely to be active during different task rule combinations than the rest of the brain, *Z* = 4.73, p = 0.0000021 (Figure 7B). The mean number of rules coded per domain was positively correlated with the percentage of activation, Spearman’s rho = 0.3754, p = 0.0000000000001719.

**Figure 7.**
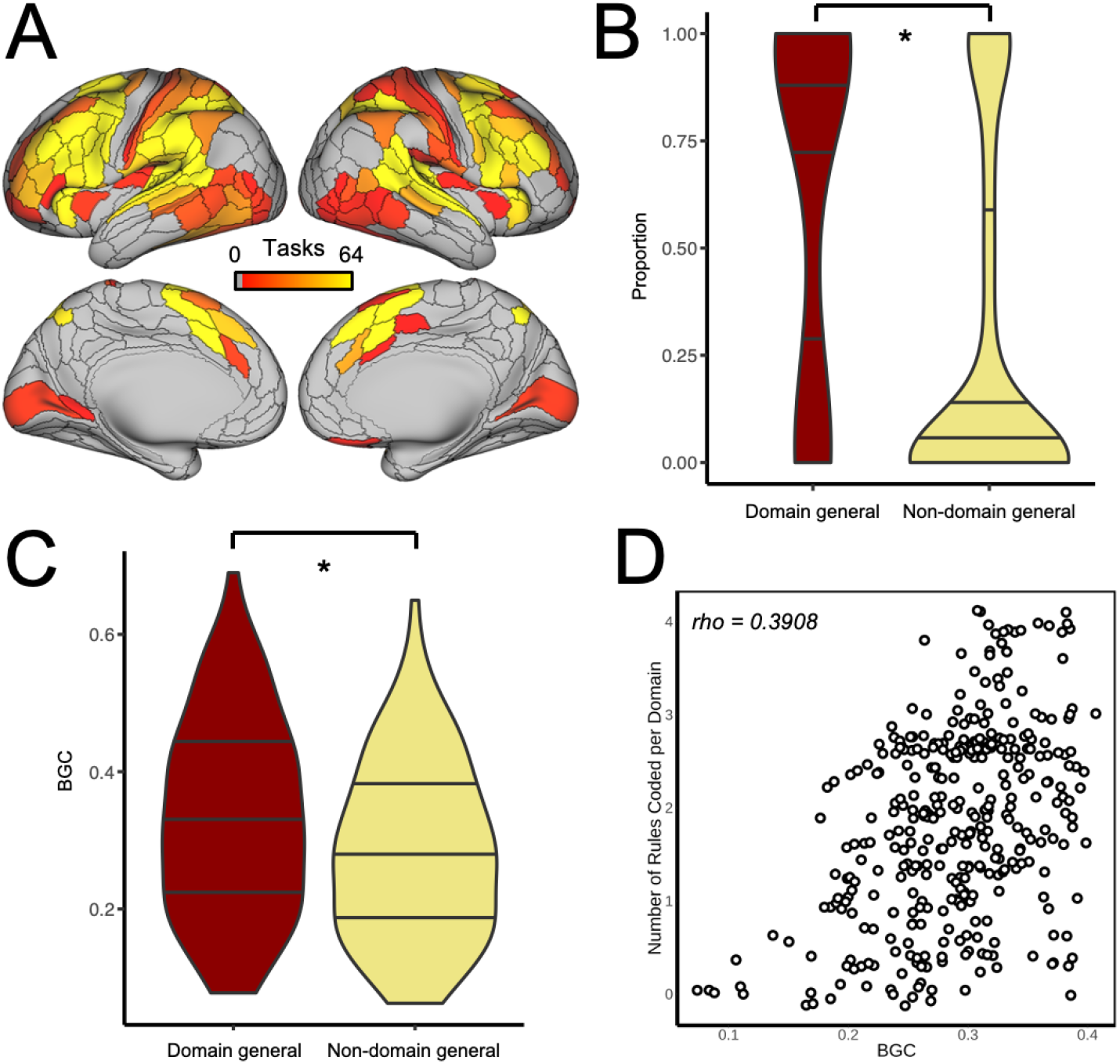
Domain general brain regions are active during a wide variety of task contexts and serve as hubs. (A) The proportion of task sets showing significant activation during the task. (B) Domain general regions, those brain areas containing information for > 3 rules per domain, have a higher probability of activation across task contexts. The lines on the violin plots represent quartiles. Asterisks indicate p < 0.05. (C) Domain general regions also have significantly greater between-network global connectivity during resting state. The lines on the violin plots represent quartiles. (D) Between-network global connectivity shows a significant positive rank correlation with their degree of domain generality.

Domain general brain regions were more likely to be activated during the performance of the task. One possible explanation for this result is that cognitive control network regions are well connected to the rest of the brain and different types of information regarding the task may converge in these areas allowing them to represent more complex aspects of the task. Using resting state data that was collected independent of task performance, we evaluated how well each brain region was connected to across-network brain regions. We used the between network global connectivity measure, which computes the average functional connectivity (weighted) of a region excluding within network connections (Ito et al., 2017; Schultz et al., 2018). We found that between network global connectivity values were significantly higher in domain general brain regions, *t*(99) = 18.99 (Figure 7C). This suggests that domain general brain regions are more likely to serve as between network hubs in the brain. Between network global connectivity was positively correlated with the mean number of rules coded per domain, Spearman’s rho = 0.3908, p = 0.00000000000001399 (Figure 6D).

### Region specific information fingerprints are predicted by connectivity fingerprints

We hypothesized that the pattern of task rule coding in a particular region would be related to the pattern of task rule coding in the brain and the degree to which those regions share information. To evaluate this hypothesis we used the concept of activity flow mapping (Cole et al., 2016) which found that activation in a held out region of the brain could be predicted by the activity in the rest of the regions of the brain weighted by the strength of the functional connections between these regions. We used a similar concept to predict information fingerprints in brain regions using the information fingerprints from other brain regions and the functional connectivity between them (Figure 8A). Here we refer to information fingerprints as the pattern of information content across all twelve of the task rules for each region of the brain. This can be visualized with a radar plot where each arm of the plot represents one of the task rules and the farther the point is from the center of the plot the more information for that rule is contained in the brain region. We also calculated the mean information fingerprint for each functional network. The somatomotor network contains the most information about the Motor rules, the visual 2 network contains the most information about the Sensory rules, and the FPN and language networks contain the most information about the Logic rules. A similar radar plot was created for predicted information fingerprints. There was a significant correlation between the network level actual information fingerprints and the predicted information fingerprints, *rho* = 0.319, *p* = 0.0001. We found that we could predict information fingerprints significantly better than chance for every region in the brain (mean correlation between predicted and actual information fingerprints of r = 0.42, Figure 8C). Regions within the same functional network by definition are most strongly connected to other regions in the same functional network. It is possible that the prediction of information fingerprints could be explained by regions within a network sharing a similar information fingerprint, and that information being weighted more heavily in our model. To address this issue, we predicted the information fingerprint for each region, but based this prediction only on information fingerprints from out of network regions. We found that we could predict information fingerprints based only on out of network sources (mean correlation between predicted and actual information fingerprints of r = 0.35). While including within network information makes for more accurate predictions, the decrease in correlation between predicted and actual information fingerprints was minimal.

**Figure 8.**
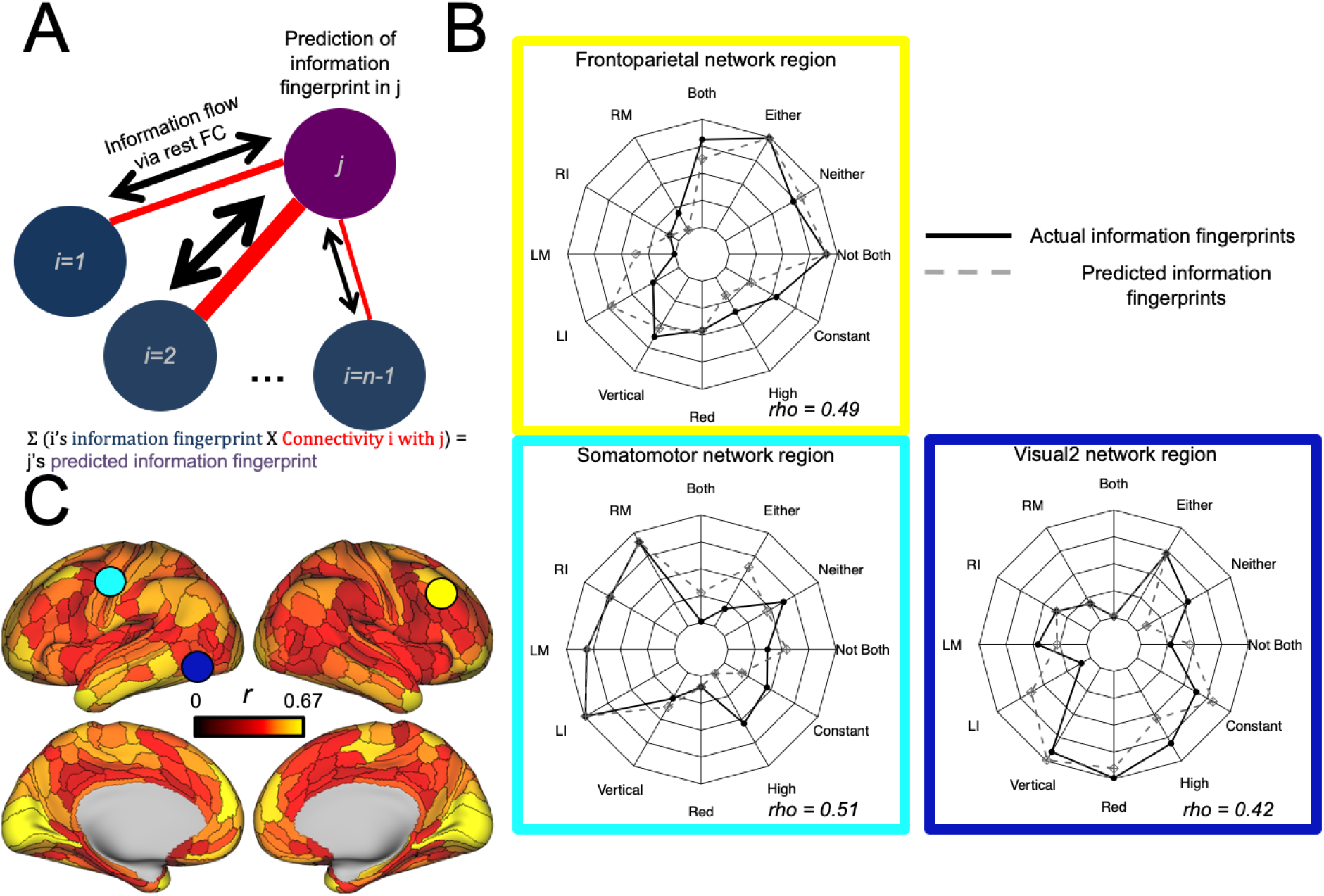
Information fingerprint mapping can predict patterns of task rule information content across the brain. (A) Information fingerprints were computed for each region of the brain. Using a simple network diffusion principle based on activity flow mapping (Cole et al., 2016), information fingerprints for a held out region can be predicted by weighting the information fingerprints from all other brain regions by the strength of their resting-state functional connectivity to the held out region. (B) Group average actual and predicted information fingerprints for three representative brain regions. Radar plots depict rank ordered information estimates with the highest information estimates at the perimeter and the lowest information estimates near the center. (C) Information fingerprints can be predicted above chance for every region in the brain. Correlations between actual and predicted information fingerprints for each brain region (FWE corrected p < 0.05). Colored circles indicate the location of the regions depicted in the radar plots in Figure 8B.

We were able to successfully predict information fingerprints. Next we tested if the domain-general brain regions we had previously identified using the actual information estimates would show a similar pattern of domain generality based on our predicted information estimates. The variability in predicted information estimates across individuals decreased relative to the variability in the actual information estimates, so – to remain conservative (reduced variability increases statistical significance) – we revised our statistical approach. We used the variability from the actual information estimates while calculating the *t* statistics for the predicted information estimates, as well as in the permutation tests. Then, similar to Figure 5A, we calculated the mean number of rules coded across domains (Figure 9A). There was a significant correlation between the predicted domain-generality map (Figure 9A) and the actual domain-generality map (Figure 5A) *rho* = 0.564, p < 0.0001 (Figure 9B). The relationship between predicted and actual rules coded for each brain region was statistically stronger than the relationship between actual rules coded and the hub properties of each region (Figure 7D), even after accounting for overlapping correlations (Meng & Rosenthal, 1992), *z* = 3.15, p = 0.0017.

**Figure 9.**
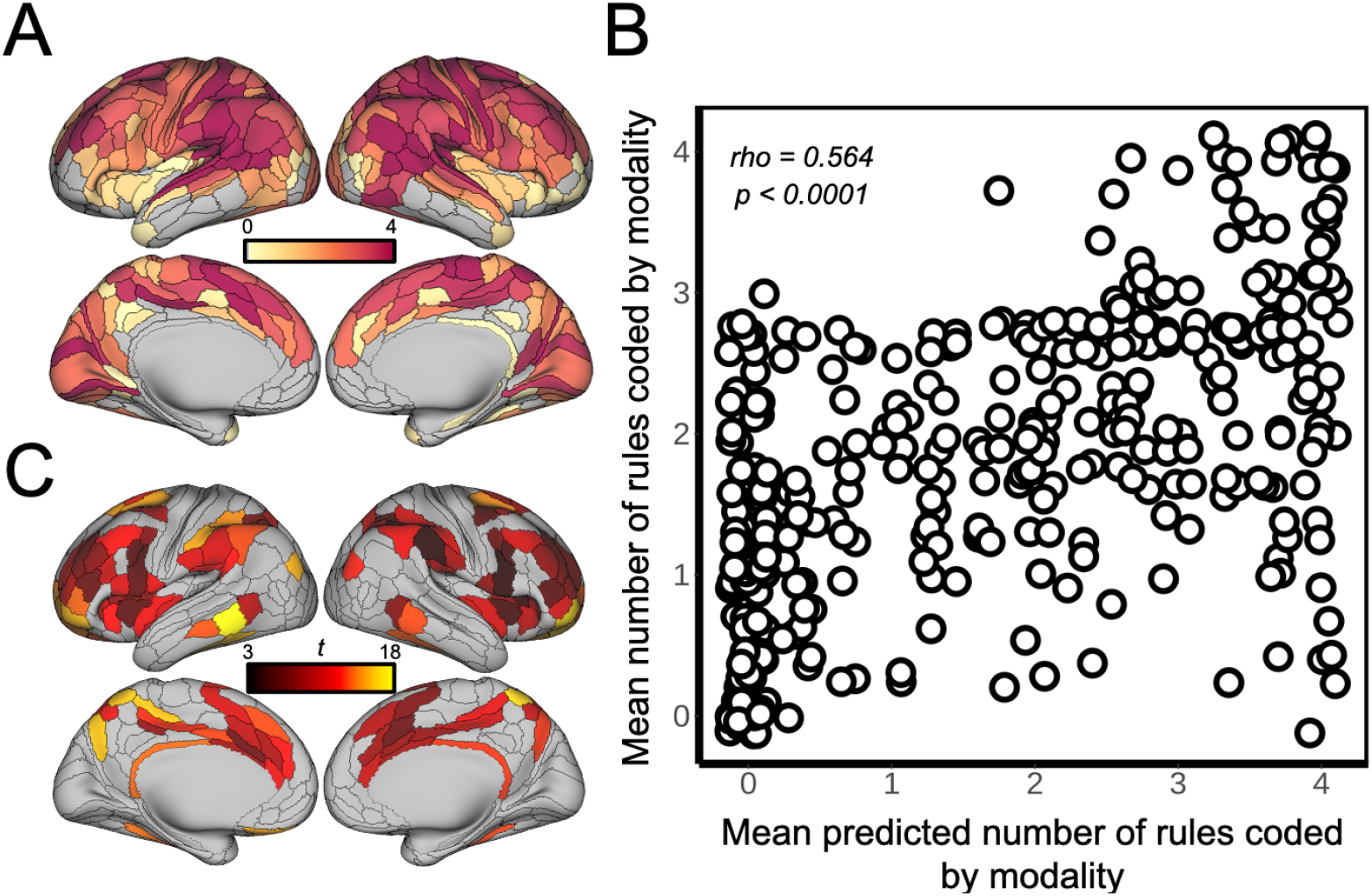
Information fingerprints and domain generality of cognitive control regions can be predicted using intrinsic functional connectivity profiles. Information estimates were predicted for each brain region by weighting information estimates in other brain regions by the strength of the connectivity between the regions (see Figure 8A). (A) Predicted information estimates for each of the task rules were thresholded to identify the mean number of rules coded for each domain based on predicted information estimates. This domain generality map for predicted information estimates was consistent with the domain generality map of the actual information estimates (Figure 5A). (B) Scatterplot depicting the significant correlation between the actual mean number of rules coded for each domain and the predicted mean number of rules coded per domain across all 360 brain regions. (C) Information fingerprints in all cognitive control regions could be successfully predicted based on the intrinsic functional connectivity patterns of those regions and the information estimates from non-cognitive control regions.

Finally, we tested the hypothesis that information fingerprints in domain-general cognitive control regions could be predicted by information fingerprints in other parts of the brain and the intrinsic functional connectivity profiles of the cognitive control regions. We recalculated predicted information fingerprints for each brain region, but based those predictions on information estimates from non-cognitive control regions. Specifically, we “lesioned” the information estimates in the CON, DAN, and FPN to see if we could predict the information fingerprints of those regions based on the information in the rest of the brain. Information fingerprints could be accurately predicted in every cognitive control region (Figure 9C). The mean correlation between the actual and predicted information fingerprints was significant for each of the three networks (CON: *M* = 0.319, *t*(99) = 21.41, p < 0.000001; DAN: *M* = 0.386, *t*(99) = 27.72, p < 0.000001; FPN: *M* = 0.374, *t*(99) = 20.41, p < 0.000001).

## Discussion

We found that task rule representations vary in how distributed they are across the cortex. While task rule representations were distributed, they tended to be represented in domain-general cognitive control networks. This is consistent with these regions being classified as hubs and showing a high degree of task activation. Furthermore, we found that the pattern of information content, or information fingerprint, for each of the rules in a brain region could be predicted using the intrinsic functional connectivity properties of each region in conjunction with the information fingerprints derived in the rest of the brain. Finally, we showed that domain generality in cognitive control networks could be predicted by the intrinsic connectivity profile of those networks in conjunction with information from other brain networks.

Previous research has suggested that task rule representation in the brain occurs somewhere on a continuum between localized and highly distributed. Consistent with this framework, we found that task rule representations varied in how distributed they were across the brain. Logic and sensory rule information was widely distributed across the brain while motor rule information was more localized to largely somatomotor network regions. We found that regions belonging to the frontoparietal, cingulo-opercular, and dorsal attention network contained domain general task rule information. These results support previous work that has identified a multiple demand network that is involved in a wide variety of cognitive tasks (Duncan & Owen, 2000). While these results support the hypothesis that task rule representation in the brain occurs on a continuum, it is important to note that this conclusion is based on the spatial limitations of fMRI data. It will be important for future studies to evaluate this hypothesis using methods capable of examining the brain on different scales.

Multivariate activation patterns have also been used to decode task rule representations. Several studies have suggested that the most accurate rule decoding using multivariate activation patterns occurs in cognitive control regions (Cole et al., 2011; Esterman et al., 2009; Reverberi, Görgen, et al., 2012; Reverberi, Gorgen, et al., 2012). If cognitive control regions can decode a larger number of different rules as we observed, it would be consistent with more accurate rule decoding in these regions as well. Electrophysiological recordings have provided additional evidence that cells in the prefrontal cortex can code for specific task rules (Asaad et al., 2000; Brincat et al., 2018; Wallis & Miller, 2003; White & Wise, 1999). The concept of mixed selectivity – the representation of multiple pieces of information by single neurons – has been used as a model to understand how cognitive control brain regions can contribute to flexible cognition (Rigotti et al., 2013). Analyses of multi-unit recordings of non-human primates have identified mixed selectivity cells in some prefrontal neurons, demonstrating that single prefrontal neurons can respond to several aspects and combinations of task variables at once (Asaad et al., 1998; Warden & Miller, 2010). This type of context-dependent neural representation is one possible mechanism for how cognitive control networks achieve their “multiple demand” status, potentially increasing cognitive flexibility via flexible representation of information.

Some brain regions exhibited domain general characteristics, but other regions coded for more domain specific information. The logic rules were preferentially represented in the default mode network. We expected that logic rules would be preferentially coded in the frontoparietal network. It is important to note that the logic rules were represented there, but that the FPN regions also contained information from the other domains which led to more balanced coding and therefore less of a preference for the logic rules. The default mode network containing information more specific to the logic domain is consistent with previous studies showing task rule decoding in the default mode network related to; the level of self-reported detail during the performance of a working memory task (Sormaz et al., 2018), representing task-related information during task switching (Crittenden et al., 2015), or covert attention during a free selection task (Haynes et al., 2007). We found that sensory networks, specifically the visual and auditory networks, preferentially coded for sensory rules. Previous research has also found evidence of coding in visual cortex for orientation (Boynton, 2005; Haynes & Rees, 2005; Kamitani & Tong, 2005) and for color (Brouwer & Heeger, 2009; Parkes et al., 2009). Multivariate patterns of activity in the auditory network can also decode auditory stimuli including human speech (Formisano et al., 2008) and decode several other categories of auditory stimuli (Staeren et al., 2009; Zhang et al., 2015). Finally, we found that the motor rules were preferentially coded in the somatomotor network. Previous studies have found that intended motor actions can be decoded from activity patterns in the motor network (Ariani et al., 2015; Gallivan et al., 2015; Gertz et al., 2017).

We observed that domain general brain regions also had a higher probability of activation during the task. This finding is consistent with descriptions of a multiple demand network (Duncan, 2010) which overlaps with cognitive control networks. We also found that domain general brain regions had higher between-network global connectivity, a measure of how well a brain region is connected to other out-of-network regions, during a resting state scan. Other studies suggest that cognitive control networks have a high degree of connectivity (Cole, Pathak, et al., 2010; Power et al., 2011), specifically for between network connections (Schultz et al., 2018). Along with a high degree of connectivity, cognitive control networks can also adaptively update these connections based on current goals (Cole, Reynolds, et al., 2013). Cognitive control networks have a high degree of connectivity and the ability to rapidly update these connections which are characteristics of flexible hubs. This is one possible explanation for why cognitive control networks show domain general characteristics.

The pattern of task rule information in each brain region could be predicted by weighting the task rule information patterns in the rest of the brain by its resting-state functional connectivity strength. Additional analyses indicated that domain generality in cognitive control networks could be predicted by their intrinsic functional connectivity profiles in conjunction with information from non-cognitive control regions. While these cognitive control networks appeared to contain domain general information, these domain general regions can be subdivided into smaller networks that specialize in different cognitive functions (Power & Petersen, 2013). The FPN is thought to be involved in the initiation of control and for adjusting control in response to feedback or instructions (Braver et al., 2003; Cocuzza et al., 2020; Cole, Reynolds, et al., 2013; Dosenbach et al., 2007). The function of the CON has been more difficult to characterize with some suggesting it has a role in task-set maintenance (Dosenbach et al., 2007; Sadaghiani & D’Esposito, 2015), arousal or alertness (Coste & Kleinschmidt, 2016), or conflict monitoring (Braem et al., 2019; Neta et al., 2014). The DAN is thought to be involved in the top-down control of attention (Buschman & Kastner, 2015; Corbetta & Shulman, 2002) and eye movement control (Corbetta et al., 1998). The heterogeneity of these components of the multiple demand network has been supported by resting-state functional connectivity data demonstrating that these components are characterized by distinct functional connectivity profiles (Ji et al., 2019; Power et al., 2011; Yeo et al., 2011). Additionally, changes in functional connectivity patterns during the performance of cognitive tasks also supports functional specialization across cognitive control networks (Cohen & D’Esposito, 2016). Other task activation studies have found evidence for domain specificity within the multiple demand network (Crittenden et al., 2016; Yeo et al., 2015), and even between portions of the FPN (Badre & D’Esposito, 2007; Nee & D’Esposito, 2016). Together, these results suggest that the domain general characteristic of cognitive control networks may emerge from the intrinsic connectivity patterns of these networks.

There are some limitations of this study which impact how generalizable the results are to other situations. One limitation is that although the C-PRO task allows us to compare many different rule combinations, the task structure is relatively rigid. Participants view instructions, are presented with stimuli, and make a response. The instructions are text presented visually. Future studies may want to consider presenting instructions with auditory stimuli, or using a non-text based visual presentation. Additionally, the stimuli used in the study were simple auditory and visual. It will be important for future studies to examine how information content is represented for other stimulus modalities, or with more complex auditory and visual stimuli. While we balanced the numerous task rules as well as possible, there were some differences in task accuracy across the rules. Ideally, participants would demonstrate equivalent accuracy on the various task rules to rule out the possibility that task accuracy was contributing to the decoding analyses we conducted. Although we cannot conclude that differences in task accuracy are not contributing to our results to some degree, we did observe largely consistent decodability across different rules within the same domain even in situations where there were task accuracy differences.

Together, our results support our hypothesis that task rule representations exist on a continuum between distribution and localization. We also found that patterns of task rule information in cognitive control networks could be predicted from patterns of task rule representation in other networks and the connectivity from those regions. This prediction used functional connectivity estimated via an independent resting state scan, which suggests that intrinsic whole-brain connectivity patterns help shape the way that task activation patterns from different brain regions are combined during task performance. Domain generality may emerge based on the intrinsic network organization of the brain. These results support the hypothesis that cognitive control networks use their unique intrinsic functional connectivity patterns to integrate domain-specific representations. These representations can then be coordinated during task performance in situations when novel combinations of rules are encountered. This likely increases the cognitive flexibility of the human brain.

## Funding

The authors acknowledge support by the US National Institutes of Health under awards K99-R00 MH096801, R01 AG055556, and R01 MH109520. The content is solely the responsibility of the authors and does not necessarily represent the official views of any of the funding agencies.

## Acknowledgements

This work was completed using the Holland Computing Center of the University of Nebraska, which receives support from the Nebraska Research Initiative. The authors acknowledge the Office of Advanced Research Computing (OARC) at Rutgers, The State University of New Jersey for providing access to the Amarel cluster and associated research computing resources. The authors also acknowledge the Rutgers University Brain Imaging Center at Rutgers University-Newark and members of the Cole Neurocognition Lab for data collection support.

## Notes

### Competing Interest Statement

The authors have declared no competing interest.

### Summary of Updates

Modifications to how the manuscript is framed; Figure 2 added for clarity; What was originally labeled Figure 8 was removed and replaced with what is now labeled Figure 9.

